# BisPin and BFAST-Gap: Mapping Bisulfite-Treated Reads

**DOI:** 10.1101/284596

**Authors:** Jacob S. Porter, Liqing Zhang

**Keywords:** bisulfite, short DNA reads, Ion Torrent, logistic, hairpin sequencing

## Abstract

**Background:** BisPin is a new multiprocess bisulfite-treated short DNA read mapper written in Python 2.7. It performs alignments using BFAST, leveraging its multithreading functionality and thorough hash-based indexing strategy. BisPin is feature rich and supports directional, nondirectional, PBAT, and hairpin construction strategies. BisPin approaches read mapping by converting the Cs to Ts and the Gs to As in both the reads and the reference genome. BisPin uses fast rescoring to disambiguate ambiguously aligned reads for a superior amount of uniquely mapped reads compared to other mappers. The performance of BisPin was evaluated on both real and simulated data in comparison to other read mappers.

BFAST-Gap is a modified version of BFAST meant for Ion Torrent reads. It uses a parameterized logistic function to determine the weights of the gap open and extension penalties based on the homopolymer run length of the DNA read. This is because the Ion Torrent sequencing technology can overcall and undercall homopolymer runs. BisPin works with both BFAST-Gap and BFAST.

BFAST-Gap is compatible with indexes built with BFAST. There are few mappers that specifically address Ion Torrent data. BFAST-Gap works with Illumina reads as well.

**Results:** BisPin with BFAST consistently had a higher amount of uniquely mapped reads compared to other mappers on real data using a variety of construction strategies. Using a hairpin validation strategy, BisPin was superior using the maximum score and mapped 73% of reads correctly.

BisPin with BFAST-Gap on Ion Torrent reads with a logistic gap open penalty function improved mapping accuracy with real and simulated data. On simulated bisulfite Ion Torrent data, the area under the curve was improved by approximately seven, and on one real data set, the uniquely mapped percent was improved by seven percent. BFAST-Gap performed better on simulated regular Ion Torrent reads than other read mappers including TMAP, designed specifically for Ion Torrent reads.

**Conclusion:** BisPin and BFAST-Gap have consistently good accuracy with a variety of data. BisPin is feature-rich. This makes BisPin and BFAST-Gap useful additions to read mapping software.

## 1 Background

Short DNA reads are treated with bisulfite to study epigenetic methylation, and these reads are mapped with software to a reference genome for epigenetic methylation discovery. Epigenetic methylation is a phenomenon where cytosine nucleic acids in DNA have a covalently bonded methyl group (CH_3_) attached to the 5 carbon of the cytosine ring. Epigenetic methylation is inheritable and plays a role in disease and development [1, 2].

Bisulfite treatment converts unmethylated cytosines into thymines, while leaving methylcytosine unchanged. The DNA sequencing process may sequence the reverse complement so that adenine corresponds to unmethylated cytosine and guanine corresponds to methylcytosine. A read mapper compares these bisulfite converted reads to a normal reference genome to discover methylcytosine, since methylcytosine should align with a cytosine in the reference genome, or in the case of the reverse complement, with a guanine. Read mapping of this sort is challenging since the bisulfite treatment introduces differences in addition to natural variation and sequencing error between the reference genome and the reads. Bisulfite read mapping has been characterized by low mapping efficiency [3], which has been shown to be correlated with reduced sequence complexity [4]. Read mapping can be time consuming with millions of short reads.

Software that maps bisulfite-treated DNA reads to a reference genome includes Bismark [5], BWA-Meth [6], Walt [7], and others. All of these programs use three phases: (1) reference genome index creation, (2) seeding, and (3) extension with alignment. A string index is usually created with either the Burrows-Wheeler transform and FM-Index or a hash table [8]. Both Bismark, which calls Bowtie2 [9] for read mapping, and BWA-Meth, which calls BWA [8] for read mapping, use the former approach. Walt uses the latter hash table approach. Seeding is accomplished by taking short subsequences of the DNA read and matching them with the string index of the reference genome. This procedure gives candidate hits, which are possible locations where the read can be mapped. Finally, an extension step is performed where the entire read is aligned to the reference genome at the candidate locations. This is normally accomplished with the Smith-Waterman local alignment algorithm. The alignment algorithm scores each location so that the best location can be reported. Differences between the reference genome and the DNA read revealed by the alignment can be either the result of genetic variation, sequencing error, or un-methylated C’s converted to T’s (or G’s to A’s on the complementary strand), and therefore, methylation at single nucleotide resolution can be determined.

Other bisulfite read mappers include BatMeth, BRAT-nova, BSMAP, BS-Seeker2, and BSmooth. BatMeth filters out reads with low entropy but does not produce output in the standard SAM file format [10]. BRAT-nova is reported to be fast but could not be made to work at the time of this writing [11]. BSMAP is outdated and cannot map reads longer than 144 base pairs, making it unsuitable for many modern sequencing projects [12]. BSMAP adds all possible combinations of C to T conversions to its index. BS-Seeker2 is similar to Bismark in that it uses Bowtie2 as a subprocess [13] for read mapping. BSmooth can call methylation marks [14]. These read mappers do nothing special to resolve ambiguously mapped reads, which are multiple high scoring alignments for a read. Their support for multiple protocols is limited to conventional sequencing protocols. They do nothing special for Ion Torrent reads.

To address these problems, this study presents BisPin, a bisulfite read mapper that deploys BFAST (BLAT-like Fast Accurate Search Tool) for read mapping. BFAST was chosen because its indexing strategy is very thorough and supports mapping with multiple indexes and spaced seeds. Spaced seeds are more sensitive [15], and indexes based on the Burrows-Wheeler transform implement a more limited spaced seed with less computational efficiency due to backtracking [16]. BFAST is feature-rich with informative output and multithreading. More importantly, BFAST has shown superior performance over tools such as Bowtie2 and BWA in the presence of indels over 10bp long [17]. This advantage becomes especially important when considering some sequencing platforms such as Ion Torrent where sequencing errors tend to introduce indels to reads [18]. Ion Torrent technology has advantages over Illumina technology such as producing longer reads [19, 20] and requiring less expensive machines [21]. Similar to Bismark and BWA-Meth, BisPin calls BFAST for read mapping, but adds special processing to accommodate short reads generated from the whole genome hairpin bisulfite construction strategy [22, 2]. A forked version of BFAST, called BFAST-Gap, was developed with special processing for Ion Torrent reads. BFAST-Gap is backwards compatible with BFAST. BFAST-Gap has an implementation of the Smith-Waterman alignment algorithm that reduces the gap open and gap extension penalties depending on the length of the homopolymer run. This should improve performance on Ion Torrent reads where gaps occur more frequently in longer homopolymer runs. BisPin enables BFAST multithreading for alignment, multiprocessing for post processing, and read partitioning for deployment on compute clusters.

## Methods

### BisPin Features

BisPin is a Python program that calls BFAST, a C++ program, to perform the three phases of read alignment and mapping. In this aspect, BisPin is similar to Bismark, which uses Bowtie2 to perform read alignment and mapping. BisPin supports directional (MethylC-Seq [23, 24]), nondirectional (BS-Seq [25]), post bisulfite adapter tagging (PBAT [26]), and hairpin [22, 2] read construction strategies. For all reads, BisPin reports a methylation calling string in the style of Bismark. The output is given as a SAM file [27]. BisPin has an alignment result summary report, which includes mapping efficiency, methylation calling statistics, timing profile information, and command line arguments. Table 1 summarizes BisPin’s features compared to several programs including Bismark, Walt, and BWA-Meth. The BisPin software is freely available at https://github.com/JacobPorter/BisPin/.

In bisulfite PCR amplification, four sequences are possible: the original forward strand, the original reverse strand, the reverse complement to the forward strand, and the reverse complement to the reverse strand. The directional construction strategy sequences the original forward and reverse strands [24]. Post-Bisulfite Adapter Tagging (PBAT) sequences the reverse complements to the original forward and reverse strands [26], and the nondirectional strategy sequences all strands [25]. Read mappers such as Bismark and Walt support mapping data with these construction strategies, and data generated with each of these methods is common. The hairpin construction strategy uses the Illumina paired-end layout to sequence the forward strand as well as the matching reverse strand with a hairpin adapter that connects the two strands [22]. The technique is especially powerful for bisulfite sequencing, as it allows for the recovery of the original untreated strand before bisulfite treatment. This is called *hairpin recovery*. This strategy was shown in previous research to improve mapping efficiency by 10% [4]. No known read mappers support special processing for the hairpin construction strategy except BisPin. BisPin has special processing for methylation calling with hairpin data, as the hairpin recovery allows the distinction between single nucleotide variants and an unmethylated cytosine.

**Table 1:** Comparison of BisPin features

| Feature | BisPin | Bismark | Walt | BWA-Meth |
| --- | --- | --- | --- | --- |
| Paired-end and single-end support | ✓ | ✓ | ✓ | ✓ |
| Directional support | ✓ | ✓ | ✓ | ✓ |
| PBAT support | ✓ | ✓ | ✓ | ✗ |
| Nondirectional | ✓ | ✓ | ✗ | ✗ |
| Hairpin support | ✓ | ✗ | ✗ | ✗ |
| Methylation calling string | ✓ | ✓ | ✗ | ✗ |
| Ambiguously mapped resolution | ✓ | ✗ | ✗ | ✗ |
| Parallelization | ✓ | ✓ | ✓ | ✓ |
| Customizable alignment scoring | ✓ | ✓ | ✗ | ✗ |
| Read file partitioning | ✓ | ✗ | ✗ | ✗ |

BisPin’s postprocessing of the BFAST raw SAM files can be done in parallel with multiprocessing, which involves six processes. Postprocessing involves choosing the highest scoring alignment for each read, rescoring ambiguously mapped reads, calling the methylation string, updating the SAM record fields, and printing a summary report.

BisPin produces a methylation calling string by matching the aligned read base to the reference base. This is done similarly to Bismark [5]. More sophisticated methods exist for methylation calling that involve read coverage and probabilistic inference [28]. For hairpin recovered data, the read sequence is compared to the recovered untreated read if available instead of the reference.

#### Rescoring Ambiguously Mapped Reads

Compared to regular short read mapping, bisulfite short read mapping produces much more ambiguously mapped reads, which are reads mapped to multiple locations [3]. Thus, it is important to have a strategy to distinguish the ambiguously mapped reads.

BisPin employs a simple and fast rescoring technique to disambiguate reads mapped to multiple locations. The rescoring technique scores different mismatches between the read and the reference differently. The matrix that represents this function can be completely specified by the user. More complicated but slower methods exist (e.g., Bayesian inference [29]) though few read mappers integrate any such disambiguation except for random assignment.The rescoring technique is a linear function of the mismatches, matches, and gaps. BisPin examines the alignment of a read to the reference genome as determined by BFAST. If the nucleotide bases match, a positive value is assigned based on the matching nucleotide base. If the bases do not match, a negative score is assigned based on the two differing bases. Gaps are scored with gap opening penalty and gap extension penalty. Different scores can be assigned for insertions and deletions. The matrix used to determine the score is the HOXD matrix in [30]. The gap open score is −400, and the gap extenstion score is −30. The gap function and the scoring matrix are the same as in Blastz [31]. The maximum scoring location, if one exists, is used to assign the read to a uniquely mapped location. A read is mapped to a unique location if there is only one location with the maximum score. This increases the uniquely mapped number of reads. The rescoring algorithm is given with Algorithm 1.

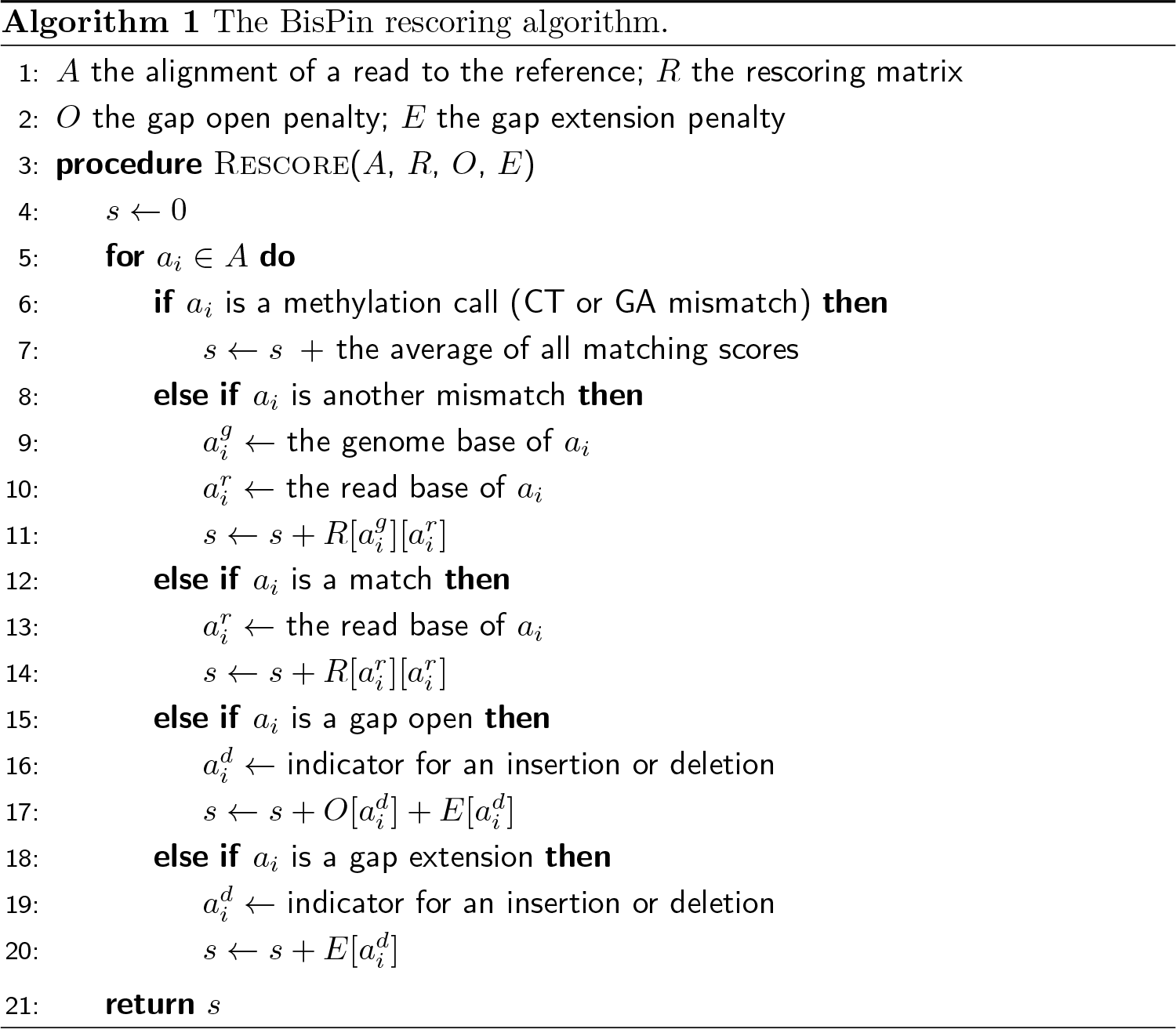

The rescoring matrix and the default BisPin alignment function are appropriate for bisuflite data and mammalian genomes. The rescoring matrix represents ratios of aligned nucleotide frequencies from noncoding mouse and human genomic regions [30]. The average matching score over all bases is used for C to T matches (alternatively, G to A matches) for the bisulfite conversion case. Bases other than C and T (G and A) should be unaffected by bisulfite treatment, so the matching and mismatching scores for those regions are the same as for regular untreated reads; thus, the functions chosen for scoring are appropriate for bisulfite data.

If the default rescoring matrix and the alignment function are inappropriate for the data, then the user can set another function. Furthermore, the exact score for the C to T (or G to A) match for bisulfite converted reads can be set with an alternative rescoring function that uses two matrices in the format given in [32]; however, the matrices in that paper were based on a uniform distribution of bases, and this does little to disambiguate ambiguous reads. In real genomes, DNA bases are not uniformly distributed [33]. On one extreme there are some Actinobacteria with GC content as high as 70% [34], and on the other extreme there is *Plasmodium falciparum* with 80% AT content [35]. The alignment function can affect the quality of results [36], so a biologically motivated scoring function, as BisPin uses, is expected to be better. Bismark appears to use an arbitrarily chosen alignment scoring function. A representation of the rescoring function is given in Figure 1.

#### Hairpin Recovery

BisPin includes special processing for the hairpin construction strategy. This data uses a hairpin connector to connect the Watson and Crick strands, and then Illumina paired-end sequencing is performed to sequence the two strands. The strands can be matched together to allow a recovery of the original strand untreated by bisulfite. This strategy is called “hairpin recovery.” A thorough description can be found in the paper [4]. For this data, BisPin recovers the original strand and uses BFAST to align it. However, not all strands will perfectly match due to sequencing error, so BisPin aligns these as regular bisulfite-treated reads either in a paired-end layout or in a single-end layout.

### BFAST-Gap Implementation

BFAST-Gap implements an adaptive weight to the gap open and gap extension penalties of the Smith-Waterman algorithm based on the length of the homopolymer run in the read. There are four function options for determining the weight: constant, logistic, exponential, and piecewise constant. The constant function is the regular method of alignment scoring that does nothing special for homopolymer runs. The exponential model opened gaps too frequently in early tests, and the piecewise constant function was thought to be too simple. For these reasons, these functions were not thoroughly examined but are provided as is. The logistic function is the most versatile, since it has portions that resemble exponential growth, exponential decay, and linear growth. Since read length is effectively bounded, the logistic function can be used to approximate exponential growth, exponential decay, and linear growth with appropriate arguments. The exponential portions of the function can approximate low order polynomials. The BFAST-Gap software is freely available at https://github.com/JacobPorter/BFAST-Gap/.

To score alignments between the read *r*_1_ and the reference *r*_2_, the following constants must be defined. Suppose that the initial gap open score is given as *g*_*o*_, and the initial gap extension score is given as *g*_*e*_. The score for two matching characters is *m*_*s*_, and the score for two mismatched characters is *m*_*n*_. The length of the homopolymer run at position *i* in read *r*_1_ is given as *D*[*i*]. The constants *g*_*o*_, *g*_*e*_, and *m*_*n*_ are negative integers, and the constant *m*_*s*_ is a positive integer.

The logistic gap open function *G*_*o*_ is a function of the homopolymer run length *D*[*i*] with slope *s*_*o*_ and center *c*_*o*_. It is given by the following:

**Figure 1:**
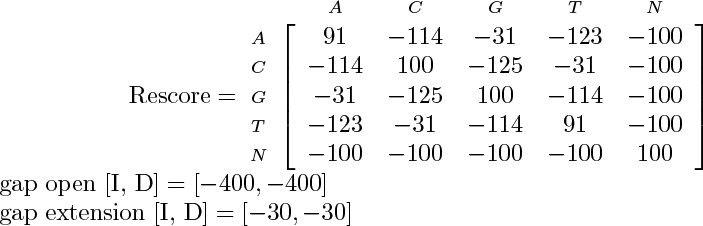
This describes the default rescoring function used to disambiguate ambiguously mapped reads. It is taken from Blastz [31, 30]. The rows of the matrix refer to the reference genome, and the columns refer to the read string. For gap opening and extension, different values can be used for insertions (I) and deletions (D). BisPin uses the average matching and mismatching scores from this matrix as default values for doing the initial alignments with BFAST.

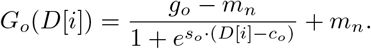

The constant gap open penalty *g*_*o*_ gives the maximum value that the function can give, and the constant *m*_*n*_, the mismatch penalty, gives the minimum value.

The logistic gap extension function *G*_*e*_, a function of the homopolymer run length *D*[*i*], has slope *s*_*e*_, center *c*_*e*_, and minimum value *z* = −1.0, and is given by the following:

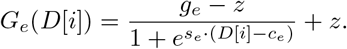

The maximum value that the function can give is *g*_*e*_, the constant gap extension penalty.

These functions are precomputed once for every execution of BFAST-Gap by storing the function values in a lookup table, since the run length parameter *D*[*i*] is discrete and effectively bounded by the maximum read length.

The run-length array is calculated once for every read by Algorithm 2. This algorithm scans through a read *r* and initially records in array *D*, at line 9, the length of a homopolymer runs ‘up to position *i*’. When a new homopolymer run is detected by detecting a change in the DNA base stored in *b*, the algorithm updates all the values in *D* associated with the homopolymer run with the total length of the homopolymer run with the **for** loop at line 11. The algorithm returns the array *D* when the outer **for** loop terminates. This algorithm has time in Θ(|*r*|), since the outer **for** loop at line 6 iterates over every base in *r* and the inner loop at line 11 iterates over every base in every homopolymer run, which comprises every base of the read.

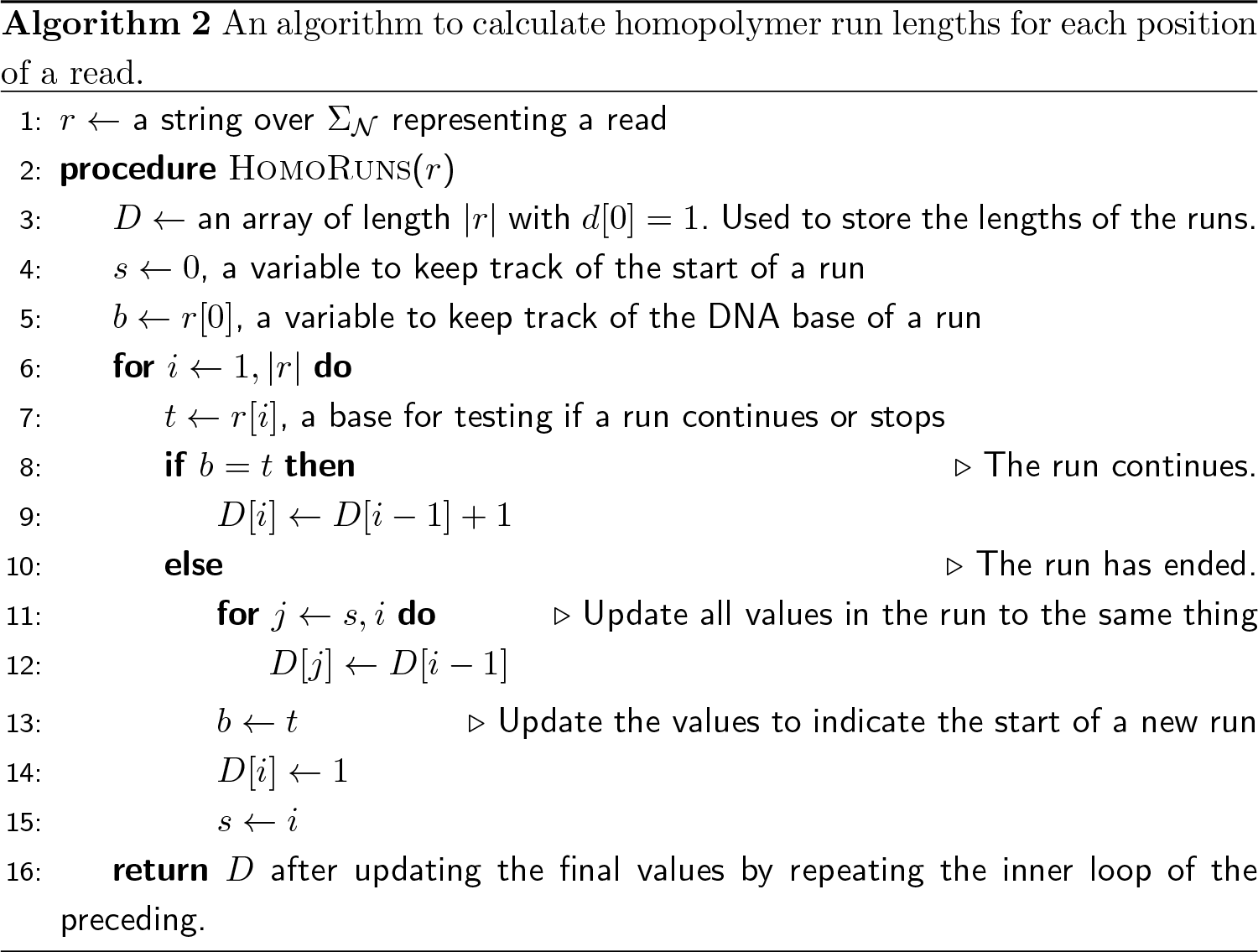

Once the run-length array *D* is computed, the alignment score and backtrace and score matrices are computed with Algorithm 3, the Run Length Smith-Waterman algorithm. This algorithm is a similar dynamic programming algorithm to the canonical Smith-Waterman algorithm except that the gap penalties are determined by the gap functions *G*_*o*_ and *G*_*e*_. The Run Length Smith-Waterman algorithm allows for negative alignment scores and an affine gap penalty. This algorithm maintains four matrices. Matrix *H* is for deletions, and matrix *V* is for insertions. Matrix *S* keeps track of the alignment score, and *B* stores the backtrace as in the canonical Smith-Waterman algorithm. The backtrace is used to compute the alignment of the read to the reference genome.

The first column and row of each matrix is initialized so that an insertion may start an alignment but not a deletion by initializing the *H* matrix row and column to negative infinity and initializing the column for the score matrix *S* and the insertion matrix *H* to the value of the gap penalty for the appropriate length of the gap. The loops at lines 21 and 22 update the interior of these matrices with the given recurrence. Finally, the position with maximum value from the bottom row is returned along with the matrices *S* and *B*. From these elements, the alignment of the entire read locally aligned to the reference genome can be computed as with the canonical Smith-Waterman algorithm using the *B* matrix. The algorithm has time complexity in Θ(|*r*_1_∥*r*_2_|), since the nested loops at lines 21 and 22 iterate over both the read *r*_1_ and the reference *r*_2_. The algorithm has space complexity in Θ(|*r*_1_∥*r*_2_|), because there are a constant number of arrays of size |*r*_1_| by |*r*_2_|.

### Data Analysis Methods

To assess BisPin (with BFAST-v0.7.0a), BisPin was compared to Bismark (v0.16.3 with bowtie2-2.2.9), BWA-Meth (with BWA-0.7.12-r1039), and Walt (v1.0) using the Genome Research Consortium’s primary assemblies of the mouse genome (GRCm38.p5) and the Arabidopsis Information Resource (TAIR) version 10 of the *A. thaliana* genome. All tests were run on Virginia Tech’s bioinformatics machine, mnemosyne2, running Red Hat Linux 4.8.5-4 with 132 GB of RAM and 16 cores comprising the Intel(R) Xeon(R) CPU E5620 @ 2.40GHz. Timing results were calculated with the Linux time command except for BisPin’s postprocessing, which used Python’s datetime module. For Ion Torrent reads only, Tabsat version 1.0.1 with TMAP version 3.4.1 was used. BFAST-Gap on regular Ion Torrent reads was compared with TMAP, Soap2 (2.21), BWA, and Bowtie2 on default settings. TMAP used the map4 algorithm.

BFAST can be run with multiple indexes that are divided into two categories: primary and secondary indexes. Whenever BFAST or BFAST-Gap were run with multiple indexes, only a single index was used as a primary index to increase run-time. Secondary indexes are used only if a match for a read cannot be found in the primary index. Unless indicated otherwise, the primary index had the mask “11111111111111111111”, and a secondary index had the mask “11111111100111111111.”

### Simulated Data

Two data sets were generated to simulate Illumina reads. Simulations were performed with Sherman v0.1.7 from Babraham Bioinformatics [37] on the mouse genome with realistic settings of error 2, CG context 20, and CHH context 98 [2]. Different read lengths were used to simulate variety in read mapping tasks. The first data set consisted of ten sets of one million paired-end 75bp reads. The second data set consisted of ten sets of one million single-end 150bp reads.

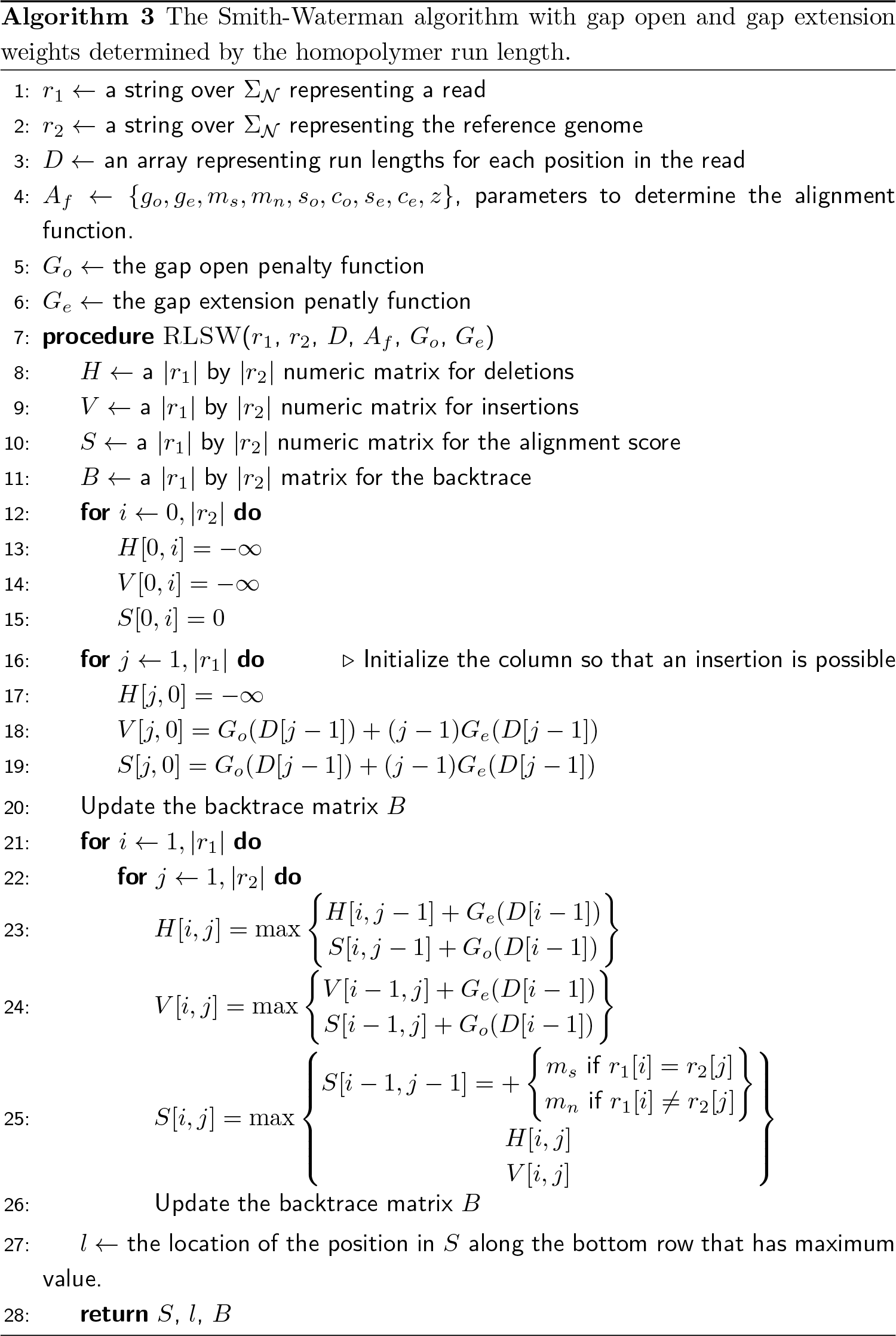

All simulated data was aligned in the same fashion as the real data. A Python script was developed to check the accuracy of read mappers. These and other Python scripts are available with the BisPin code. Only uniquely mapped reads were checked and considered accurately mapped if the starting location was within three bases of the real location. Average precision and recall were computed. Precision is the ratio of the number of correctly uniquely-mapped reads and the number of uniquely mapped reads. Recall is the proportion of the uniquely mapped reads that are correctly mapped of the total number of reads.

A test to compare the rescoring function against a random choice was performed to see if the rescored read was more likely to align correctly than random chance. This test used 100,000 reads from the 150bp single-end simulated data. The set of rescored ambiguously mapped reads was extracted, and for each read, a random alignment location was chosen. The percentage of these that were correctly mapped was compared to the percentage of the rescored reads that were correctly mapped. This process was repeated 32 times to calculate a *p*-value.

The preceding Sherman simulated data was used to test BisPin on simulated bisulfite-treated Illumina reads. BisPin with BFAST and BFAST-Gap were tested on simulated Ion Torrent reads generated by DWGSIM since it simulates the undercalling and overcalling of homopolymer runs found in Ion Torrent reads [38]. For DWGSIM, the bisulfite treatment was simulated on the reference genome with the CpG rate as 0.215, the CH rate 0.995, and the over conversion and under conversion rates of 0.0025 [3, 2]. This was done with custom Python scripts provided by Hong Tran. Custom Python scripts were used to simulate single-end data from DWGSIM. DWGSIM was used with the following realistic settings: “dwgsim -e 0.012 -E 0.012 -d 250 -s 30 -S 0 -N 1000000 -c 2 -1 200 -2 200 -f TACGTACGTCTGAGCATC-GATCGATGTACAGC” [39].

These settings were used to produce two data sets, one data set for simulated bisulfite-treated reads, and the other data set for regular Ion Torrent reads. The simulated bisulfite-treated Ion Torrent reads were divided into a 10k read training set and a one million read test set in order to tune the logistic gap penalty functions.

The logistic gap open function for BFAST-Gap was trained on the 10k read training set by looking at slopes from 1.2 to 0.5 and centers from −20 to 150. A single index was used with mask “11111111100111111111.” The alignment function had values of 96 for a match, −90 for a mismatch, −600 for a gap open penalty, and −50 for a gap extension penalty. The optimal slope (*s*_*o*_ = 1.0) and center (*c*_*o*_ = −15) were chosen based on which values maximized the F1-score and the area under the curve for all BisPin filter values from 0 to 96.

Once the logistic gap open function was tuned, the settings for the logistic gap open function were set, and the logistic gap extension function was tuned in a similar manner. This resulted in a slope of *s*_*e*_ = 0.1 and a center of *c*_*e*_ = 90. The tuned logistic gap open and extension functions are considered the default settings for BFAST-Gap. Figure 2 gives the tuned logistic gap extension penalty function. This function is approximately constant until a run length of approximately 50 base pairs.

The entire one million test set was mapped with BisPin calling BFAST-Gap with these settings. One execution was run with constant open and extension penalties, another execution was run with the logistic open penalty function and a constant extension penalty function, and a third execution was run with logistic open and extension penalty functions with the tuned parameters. The F1-score was calculated such that an alignment was considered correct if it was within three bases of the correct location on the reference genome. BisPin used two indexes, and rescoring for Ion Torrent reads was turned off since the floating point logistic function tends to resolve ambiguously mapped reads.

**Figure 2:**
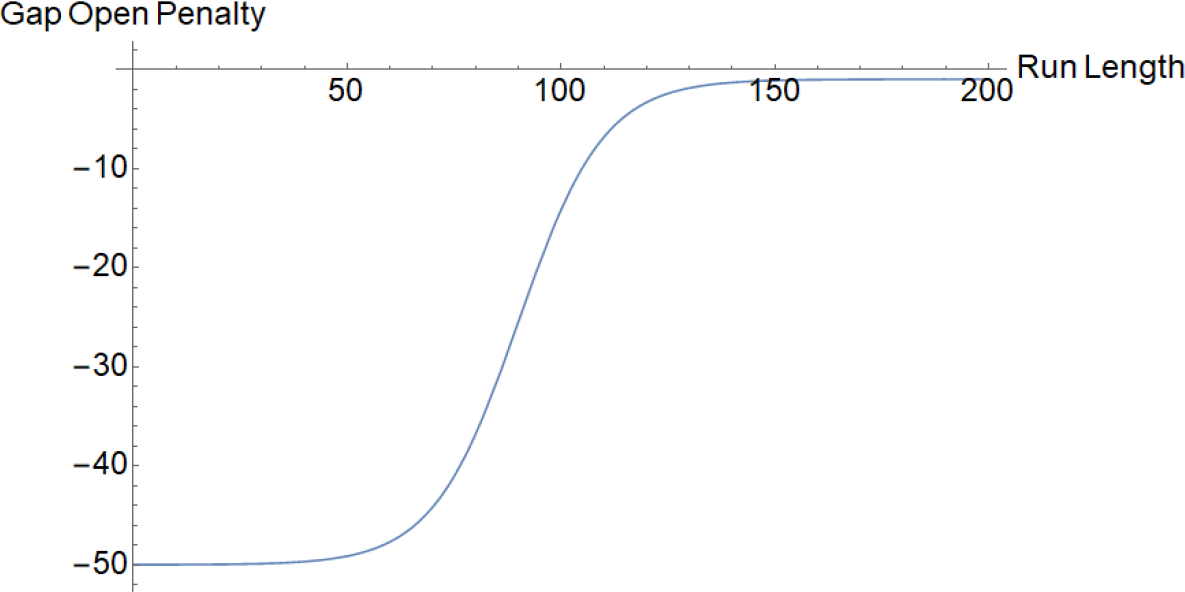
An example of a logistic function for the gap extension penalty for BFAST-Gap. This function has *g*_*e*_ − *z* = −49, *m*_*n*_ = −50, *s*_*e*_ = 0.1, and *c*_*e*_ = 90.

For one million DWGSIM simulated regular Ion Torrent 200bp reads, BFAST-Gap was run with the tuned logistic gap open and logistic gap extension penalty functions without a filter and two indexes. TMAP did not have a filter and was run with default settings for the map4 algorithm. Bowtie2, BWA, and Soap2 were run with default settings. Soap2 could not be accuracy tested since it does not produce a SAM file. For the simulated regular Ion Torrent data, precision and recall were calculated with the same Python script as other simulations.

### Real Data

Variegated real data was downloaded from the SRA (Sequence Read Archive) [40]. Each Illumina data set had reads of approximately 100bp length. The mouse data used 100k reads, and the plant data used 500k reads. Only SRR4295457 data was paired-end. The rest were single-end. Hairpin 101bp mouse data was constructed with the procedure given in [2]. Table 2 gives a summary of the real Illumina read data used in this study. The mapping efficiency was calculated with each mapper’s self report except for BWA-Meth since it did not give such a report. For BWA-Meth, a Python script was created to determine the mapping efficiency. If a read had only one location, it was uniquely mapped. If it had several locations including locations given in the **XA:Z** tag used by BWA, the read was classified as ambiguously mapped. If the alignment had the unmapped or filtered flag set, it was classified as unmapped. The default settings of all mappers were used.

Different data sets came from different sequencing construction strategies, which included directional, nondirectional, and PBAT constructions as indicated in Table 2. If a read mapper did not support the sequencing construction strategy, the read mapper was not run on that data set. Support for these construction strategies is indicated in the read mapper’s interface.

BisPin used one primary index and two secondary indexes for mouse data but only one secondary index for A. *thaliana* data. These indexes work such that if a candidate location cannot be found in the primary index, then the secondary indexes are tried. The primary indexes searched for a seed with a length 20 exact match, and the secondary indexes used spaced seeds. Rescoring was on.

**Table 2:**
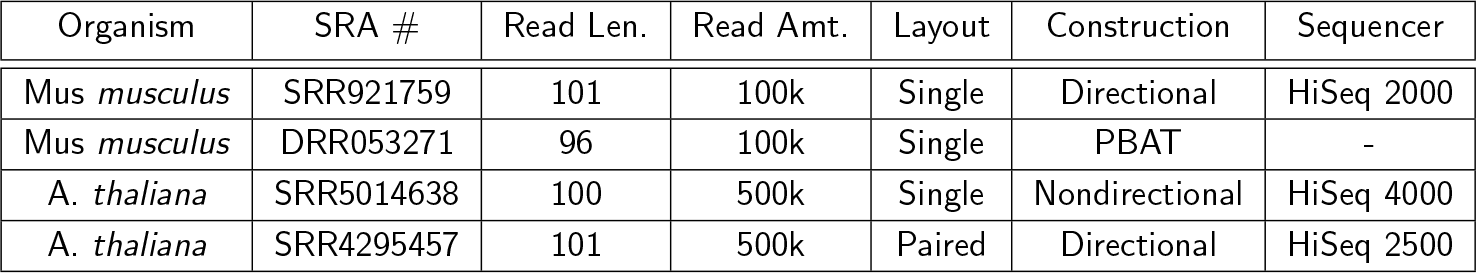
A summary of the real Illumina read data features. All data was obtained from the Sequence Read Archive (SRA) at https://www.ncbi.nlm.nih.gov/sra.

| Organism | SRA # | Read Len. | Read Amt. | Layout | Construction | Sequencer |
| --- | --- | --- | --- | --- | --- | --- |
| *Mus musculus* | SRR921759 | 101 | 100k | Single | Directional | HiSeq 2000 |
| *Mus musculus* | DRR053271 | 96 | 100k | Single | PBAT | - |
| *A. thaliana* | SRR5014638 | 100 | 500k | Single | Nondirectional | HiSeq 4000 |
| *A. thaliana* | SRR4295457 | 101 | 500k | Paired | Directional | HiSeq 2500 |

To test the logistic gap open function, the optimal slope and center from the simulated data was used, and BisPin with BFAST-Gap was run on three real data sets without an alignment quality filter using both a constant and a logistic gap open penalty function. The quality filter for BisPin was turned off since the logistic gap open function changes the alignment score making the alignment scores between the two alignments incomparable. The amount of uniquely mapped reads between these data sets indicates which alignment scoring function performs better. The real bisulfite-treated Ion Torrent data was downloaded from the SRA trace archive, with two mouse datasets (SRR1534391 and SRR1534392) containing one million reads each, and one human dataset (SRR3305017) containing 251,374 reads. The data was aligned with one index when testing BisPin with BFAST-Gap and was aligned with two indexes when testing BisPin with BFAST. Rescoring was turned off.

Real untreated mouse and human data were used to test BFAST-Gap with tuned logistic gap open and extension penalty functions on regular Ion Torrent reads. The SRA numbers were mouse ERR699568, human SRR2734774, and human SRR611141. Each data set had one million reads. The regular mapper programs TMAP, Bowtie2, BWA, and Soap2 were used for comparison. The uniquely mapped, ambiguously mapped, and filtered/unmapped percentages were calculated based on the programs self-report for Bowtie2 and Soap2. Custom Python scripts available with the BisPin source code were used to calculate these statistics for BFAST-Gap, TMAP, and BWA. BFAST-Gap was run with a stricter filter of 75 on its default settings with two indexes, and TMAP was run with a filter of 0.625, which approximately corresponds with a BFAST-Gap filter of 60. Other read mappers were run on their default settings.

## Results

### Simulated Data

BisPin had the highest recall for the single-end data and the highest precision for the paired-end data as shown in Figure 3. Both BisPin and Bismark correctly called the approximate amount of methylation in each context, 80% for CpG and 2% otherwise.

Interestingly, the recall generally decreased by approximately 0.10 for the one million paired-end 75bp reads for all the read mappers except for BWA-Meth. The F1-score, a balanced average of precision and recall, for this data was the following, 0.79 (BisPin), 0.80 (Bismark), 0.91 (BWA-Meth), and 0.75 (Walt).

**Figure 3:**
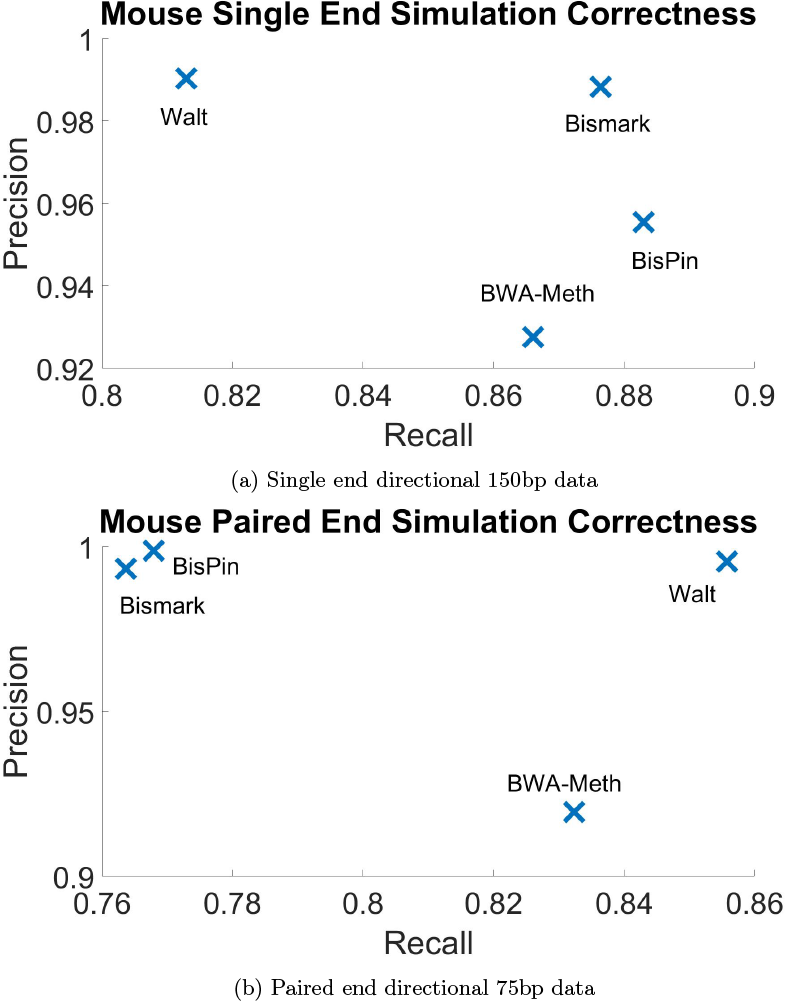
The average of precision and recall over ten replicates of one million read; of uniquely mapped reads on Sherman mouse read simulations with error 2, C conversion 20, and CH conversion 98. The standard deviation for all statistics for all data was below 0.0006.

Data consisting of 32 replicates of 100k 150bp single-end reads was used to assess the rescoring functionality. Choosing a random alignment for a read from the set of rescored reads mapped 19.1% correctly on average while the rescoring functionality gave an average of 23.5% correct alignments. The rescoring approach always returned more correctly aligned reads than the random choice with an average difference of 4.4%. This suggests a *p*-value close to zero and that there is a statistically significant difference in these distributions such that the rescoring functionality is better than random assignment for uniquely aligning some reads.

Using BisPin with BFAST on default settings and with BFAST-Gap on tuned gap penalty functions, the one million DWGSIM reads were mapped, and precision, recall, and F1-score were calculated as shown in Figure 4. BWA-Meth and Bismark had moderate performance, while Walt performed poorly. These read mappers appear ill-suited for mapping Ion Torrent reads. BisPin and Tabsat performed the best. BisPin with BFAST-Gap had slightly higher precision and slightly lower recall but with a higher F1-score than BisPin with BFAST.

**Figure 4:**
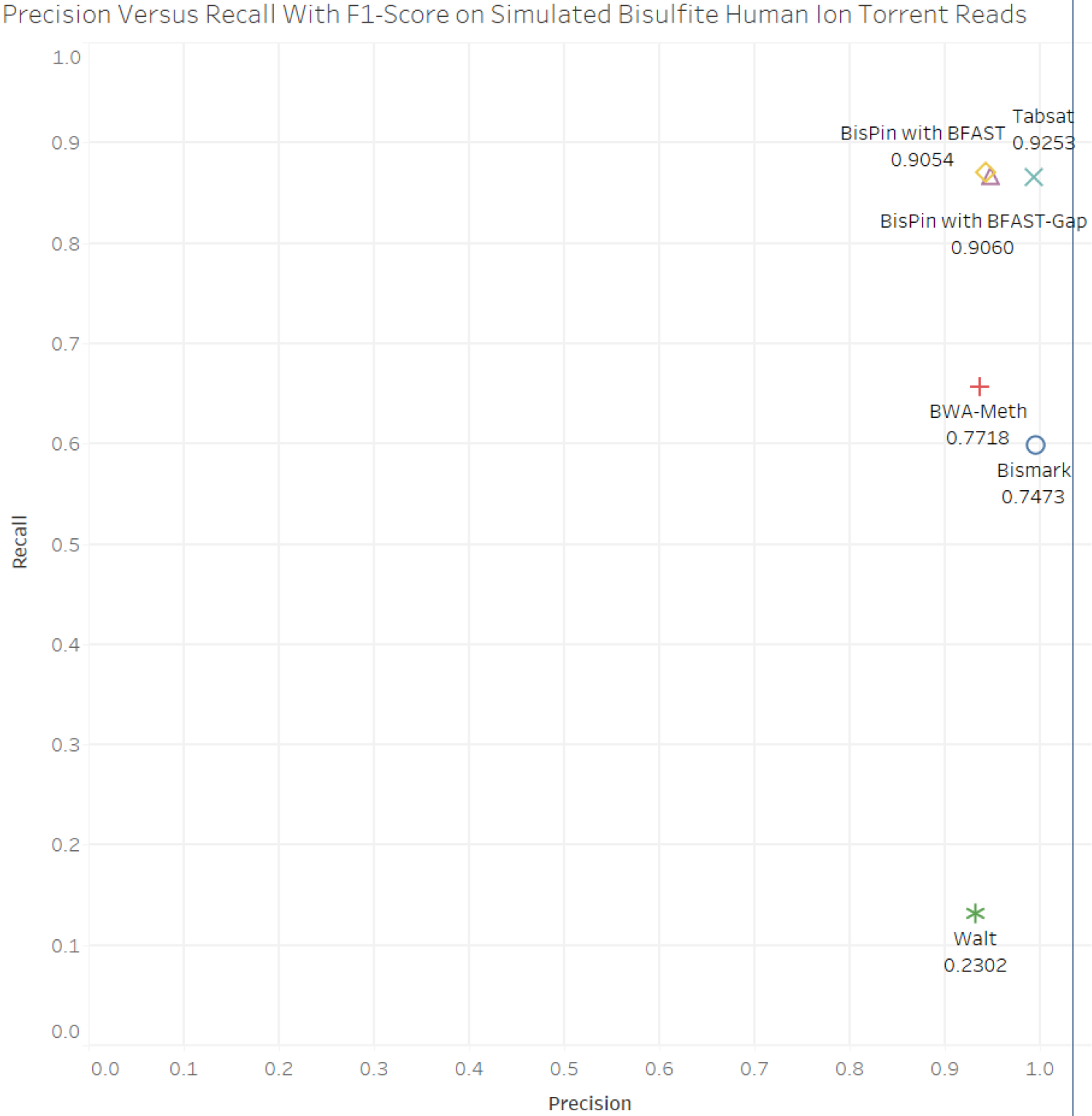
Precision versus recall on simulated human bisulfite-treated Ion Torrent reads. The numeric label is the F1-score.

The proportion of reads in each mapping category, uniquely mapped, ambiguously mapped, and unmapped or filtered, on the same simulated data for each read mapper is shown in Figure 5. BisPin and Tabsat performed the best with BisPin having about five percent more reads uniquely mapped with few ambiguously mapped. Bismark and BWA-Meth were less good, and Walt performed poorly.

**Figure 5:**
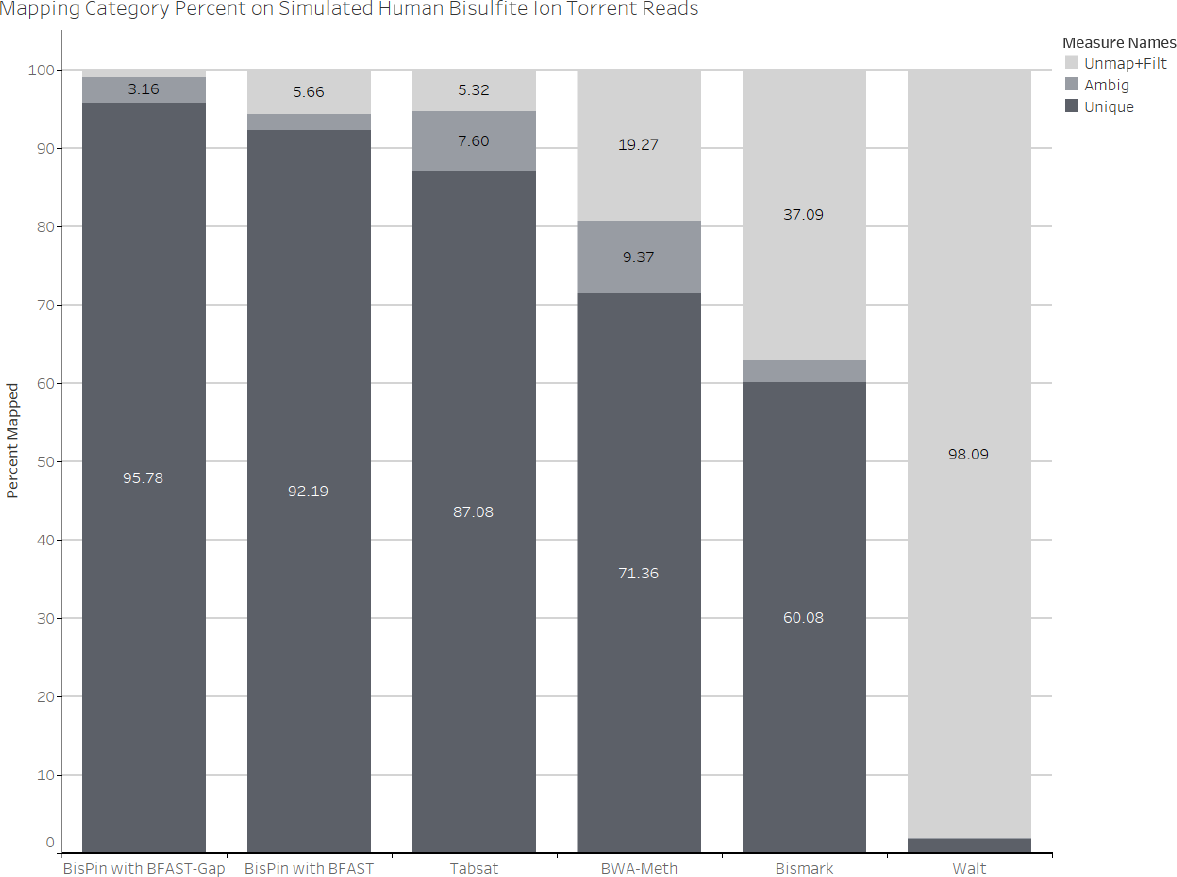
Mapping performance by category on one million DWGSIM simulate bisulfite Ion Torrent reads.

Tuning the logistic gap open function on BFAST-Gap on 10k simulated Ion Torrent reads revealed interesting results, as shown in Figure 6. Settings with low slopes tended to perform the worst with low centers performing the best among these. Settings with slopes closer to 1.0 performed the best with centers around 0 to −30; however, with centers much larger than this, performance was worse. From this analysis, a slope of 1.0 and a center of −15.0 was chosen.

After tuning, BisPin with BFAST-Gap was run on the one million simulated reads test set, and the alignment with the tuned logistic gap open function achieved superior area under the curve and F1-score, as can be seen in Figure 7. Executions with two indexes and one index were performed, and using two indexes noticeably improved the performance. The maximizing filter value for the constant penalty function was 65, and for the executions involving the logistic function, it was 85. The logistic function tends to reduce alignment scores, so a higher filter value is needed to compensate. The AUC was improved by approximately 7, for the two index case, and the F1-score was improved by 0.003. This indicates that using the logistic gap open function and a constant gap extension penalty function improved mapper performance on this simulation. The executions where both the gap open and extension functions were tuned logistic functions performed nearly identically to the executions with the logistic gap open penalty function and a constant gap extension function but with slightly worse performance.

Figure 8 shows the precision, recall, and F_0.5_-score for mappers on the simulated DWGSIM regular Ion Torrent data. BFAST-Gap had the highest overall score. TMAP performed the worst, which is surprising since it is designed for Ion Torrent reads.

**Figure 6:**
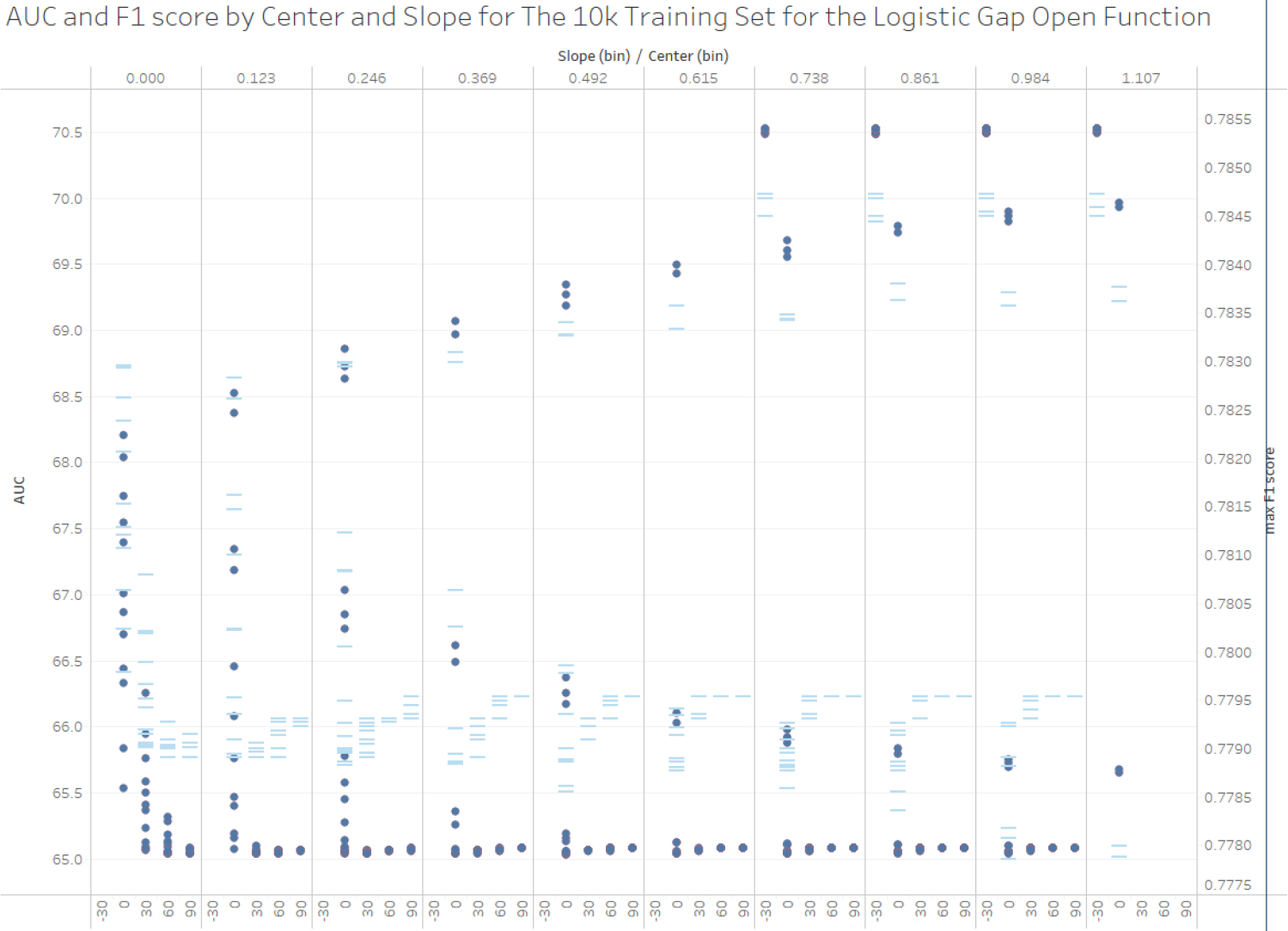
AUC and F1-Score by Slope and Center for the Logistic Gap Open Penalty Function on 10k Simulated BS Ion Torrent Reads. The circles represent AUC scores and the bars represent maximum F1-scores for alignment quality filter values ranging from 0 to 96.

### Real Data

The percentage of uniquely mapped reads gives a reasonable test of read mapper performance, and recent papers use this method [11, 7]. The simulated data shows that the percentage of uniquely mapped reads that are correctly mapped is very high. Although the Sherman Illumina simulated data does not include indels, indels are an order of magnitude more rare than SNPs, so they should not affect mapper performance as much [41]. This suggests that many of the uniquely mapped reads are probably aligned correctly.

Figure 9 summarizes the mapping efficiency of the read mappers on the four real data sets. In all cases, BisPin had the most uniquely mapped reads. Without rescoring, BisPin would have reported approximately 10% ambiguously mapped reads, but the rescoring moved from 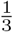 to 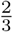 of these to uniquely mapped. BisPin and Bismark reported on the percentage of methylation in all contexts.

On 100k reads of hairpin mouse embryonic data, BisPin did hairpin recovery on 62% of the reads. BisPin mapped this data as regular paired-end data with 81.7% uniquely mapped, but other read mappers mapped few reads uniquely in paired-end mode. This is probably because paired-end mapping usually assumes that one end is upstream of another rather than overlapping as with hairpin data.

A method of validating the correctness of the read mappers was performed with the hairpin data. One million reads were sampled, and 604431 reads were hairpin recovered. These reads were mapped with BFAST, with BisPin’s defaults, and with Bowtie2 and BWA on their default settings. BisPin, Bismark, and BWA-Meth were run on the one million bisulfite reads, and the uniquely mapped reads were compared with the uniquely mapped (hairpin recovered) original reads from each regular read mapper. If the starting location of the bisulfite read was within three bases of the mapped original read, then it was considered correctly mapped. The motivation for this is that the original read should more often map correctly because bisulfite treatment has a tendency to reduce sequence complexity, which can adversely affect mapping quality [4]. These results are summarized in Table 3. BisPin did well with the highest correct when compared to each regular read mapper

**Figure 7:**
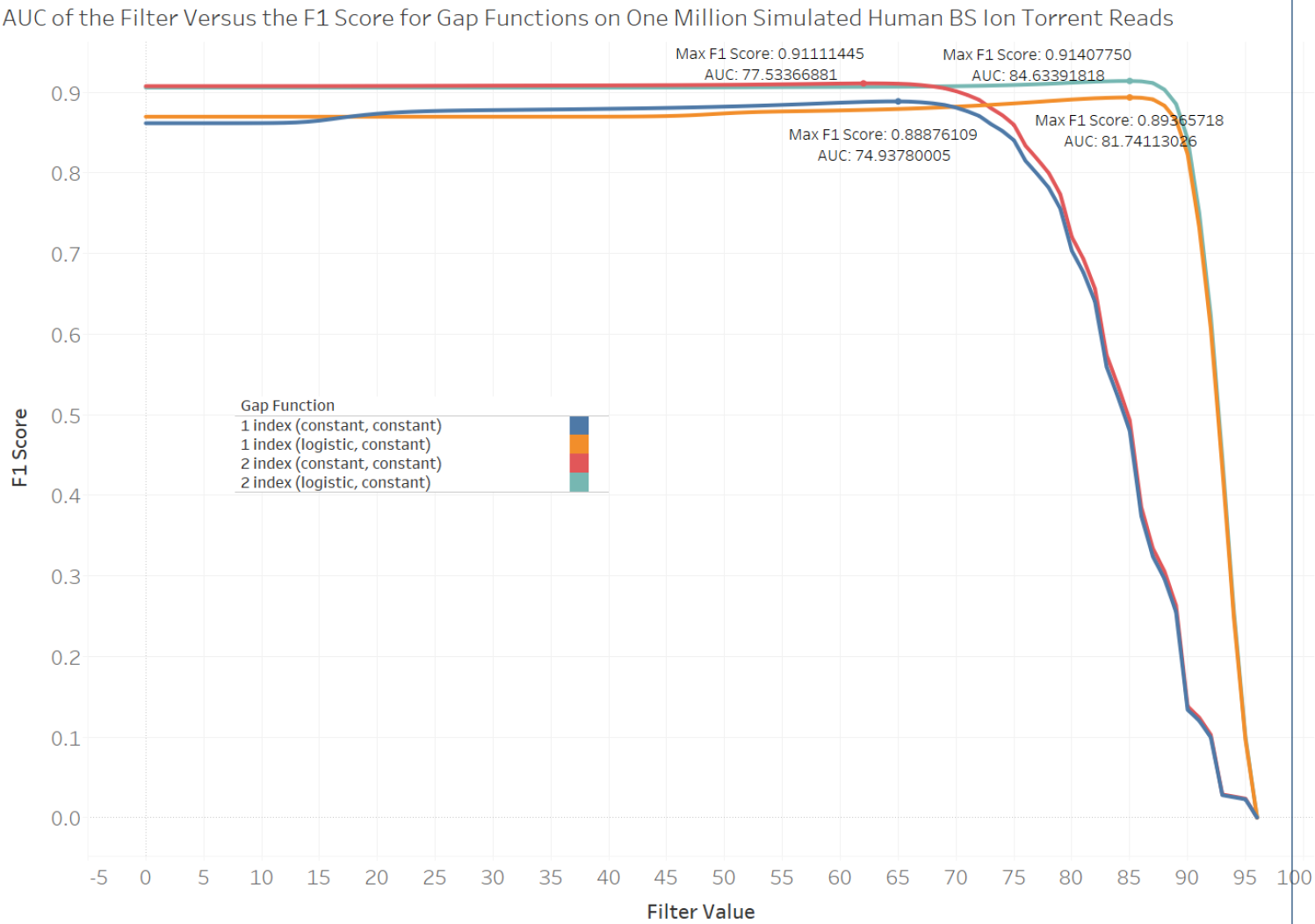
Performance of BisPin with the tuned BFAST-Gap logistic gap ope function on one million simulated test reads as measured by area under the curv and F1-score using one index and two indexes.

**Table 3:** Hairpin validation. BFAST, Bowtie2, and BWA were used to map the 604,431 original recovered reads from one million bisulfite reads, and the percent of these recovered reads that were uniquely mapped to the same location with each bisulfite read mapper on the one million bisulfite reads was reported. The maximum value is in bold, and BisPin had the highest value. Walt had the highest average, but BisPin was second with a difference of 0.8%.

| Bisulfite Mapper | BFAST | Bowtie2 | BWA | Average |
| --- | --- | --- | --- | --- |
| BisPin | **73.8161%** | 60.5174% | 60.1448% | 64.8261% |
| Bismark | **63.3881%** | 53.0212% | 53.8516% | 56.75363% |
| BWA-Meth | **66.3428%** | 56.926% | 61.8104% | 61.693067% |
| Walt | **72.2992%** | 62.6082% | 62.0344% | 65.647267% |

**Figure 8:**
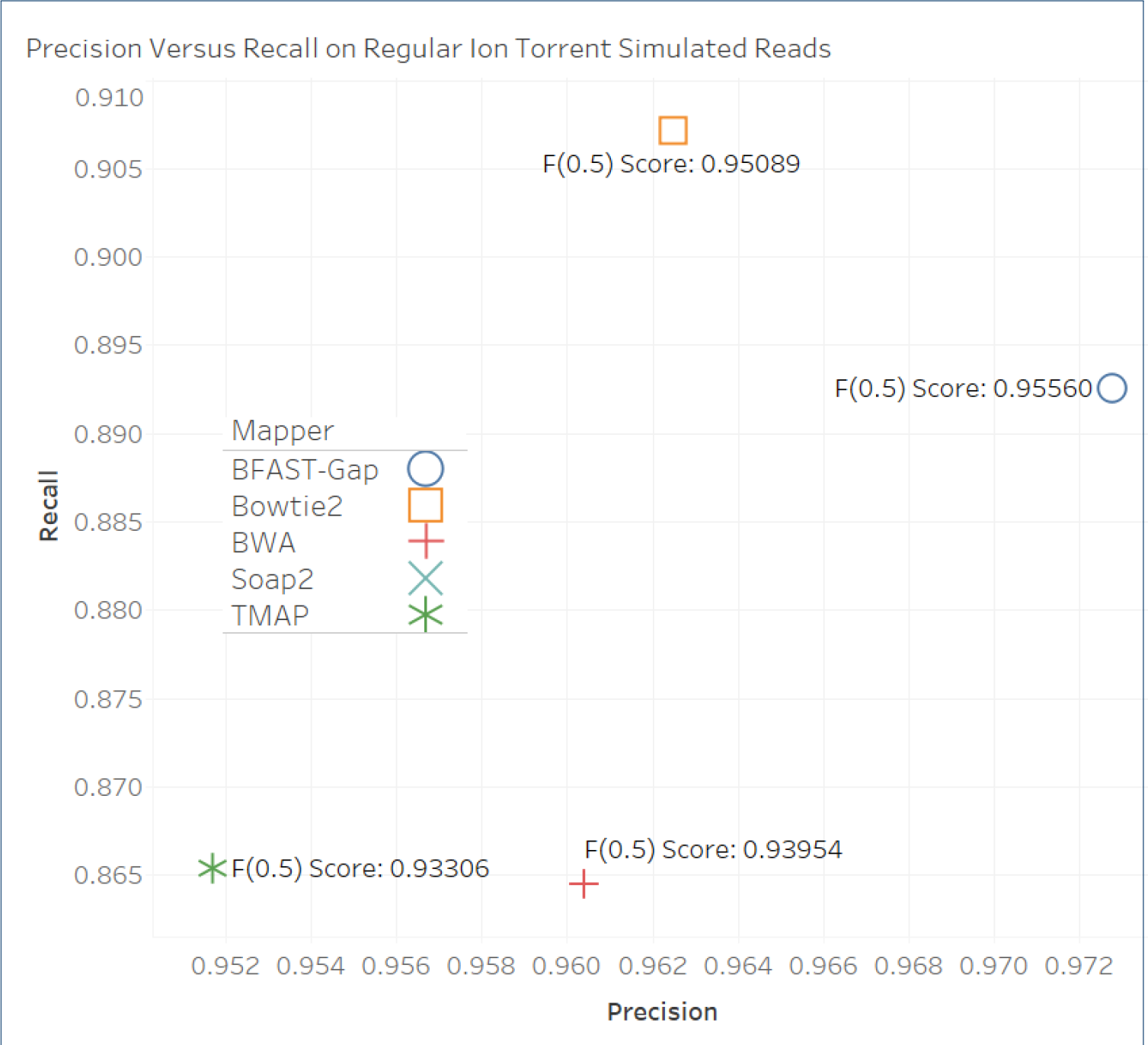
The precision, recall, and F_0.5_-score for regular Ion Torrent DWGSIM 1 million simulated 200bp single-end reads with settings e=1.2 and E=1.2. BFAST Gap has the best F_0.5_ score and the highest precision. Default settings are used throughout. Soap2 is not shown since it did not produce SAM file output.

To test the performance of BisPin with BFAST on bisulfite Ion Torrent data, it was run on default settings along with other mappers with performance results indicated in Table 4. BisPin had the most uniquely mapped for all five data sets, and Tabsat was the second best. For the mouse data, BisPin mapped more than ten percent of the reads uniquely compared to Tabsat. Walt was the worst with very poor performance.

The read mappers were run with their default settings because that is what users are likely to do. Bismark, BWA-Meth, and Walt perform poorly on the bisulfite-treated Ion Torrent reads in Table 4 because they may not be tuned to the read lengths and error profile of Ion Torrent data. To test if improvements could be made, tuning parameters were adjusted and the read mappers were run again on the mouse SRR1534391 data. BWA-Meth offers no tuning parameters, so there was no attempt at improving its performance. Bismark has a minimum score function that filters out low quality alignments. This was changed to ‘L,0,−0.8’ and this produced 76.6% uniquely mapped reads, over 50% of an improvement. Walt has a maximum number of mismatches parameter with a default of 6, and increasing this to 175 increased the uniquely mapped percent to 17.31%, only approximately 13% of an improvement. This was the only parameter that could be tuned to make an improvement. Thus, Bismark’s performance can be greatly improved, but Walt’s performance improvement seems limited for Ion Torrent data. Walt filters out reads less than 38 base pairs in length, and this was 8.56% of the SRR1534391 data. This makes BWA-Meth and Walt less suitable for Ion Torrent data.

**Figure 9:**
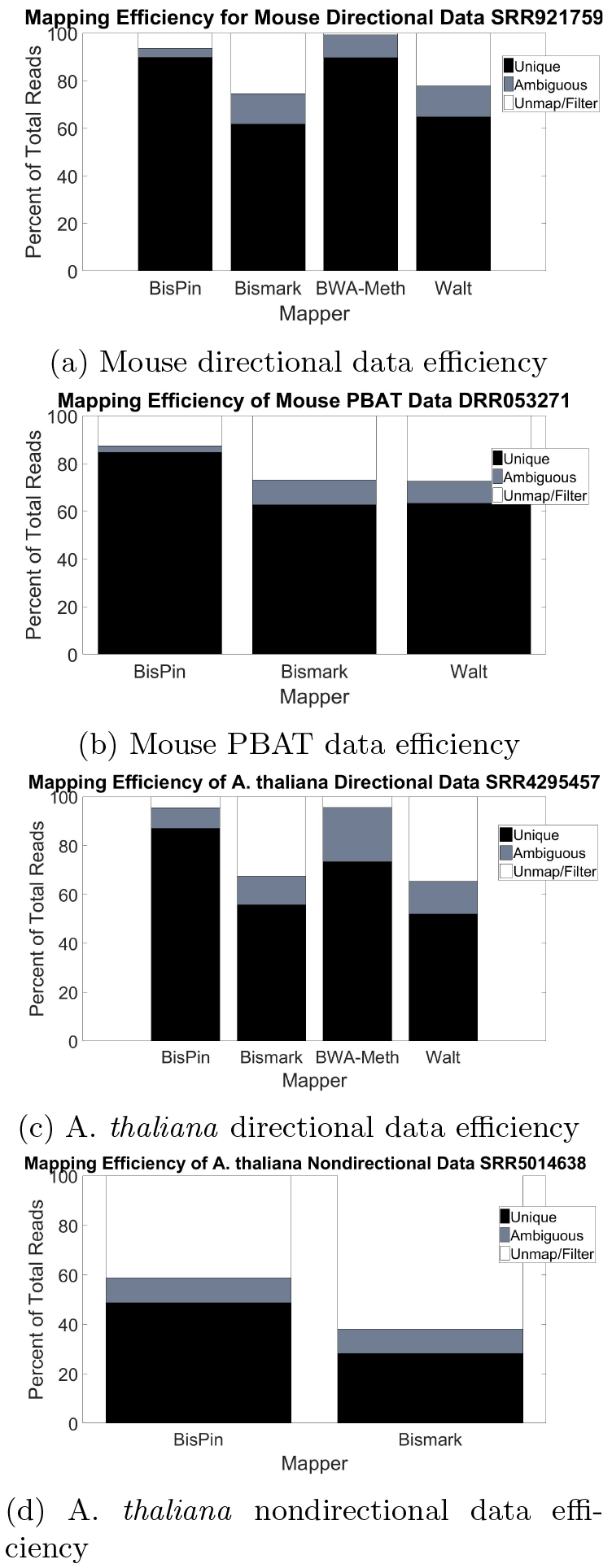
BisPin mapping efficiency on real data compared to other mappers. BisPin always has the highest number of reads uniquely mapped. Figure 9c is paired-end data, but all other data is single-end.

Since Ion Torrent reads can vary in length, a test of mapper performance in relation to read length from the SRR2842547 and SRR2842546 data was performed.

**Table 4:** The percent of real bisulfite-treated Ion Torrent reads uniquely mapped on five data sets. BisPin performs the best.

| SRA Mouse # | BisPin | Bismark | BWA-Meth | TabSAT | Walt |
| --- | --- | --- | --- | --- | --- |
| SRR1534391 | 91.79 | 21.20 | 45.17 | 80.65 | 3.79 |
| SRR1534392 | 91.38 | 16.60 | 38.79 | 80.05 | 3.77 |

| SRA Human # | BisPin | Bismark | BWA-Meth | TabSAT | Walt |
| --- | --- | --- | --- | --- | --- |
| SRR2842546 | 90.00 | 60.60 | 70.42 | 84.43 | 19.19 |
| SRR2842547 | 89.33 | 60.70 | 68.28 | 83.93 | 18.34 |
| SRR3305017 | 92.63 | 35.70 | 65.32 | 86.67 | 23.26 |

Figure 10 gives a histogram of read lengths from the SRR1534392 data. This distribution has a long tail with a significant mode at the high end of the distribution indicating that many reads are quite long, but a substantial amount have short lengths.

**Figure 10:**
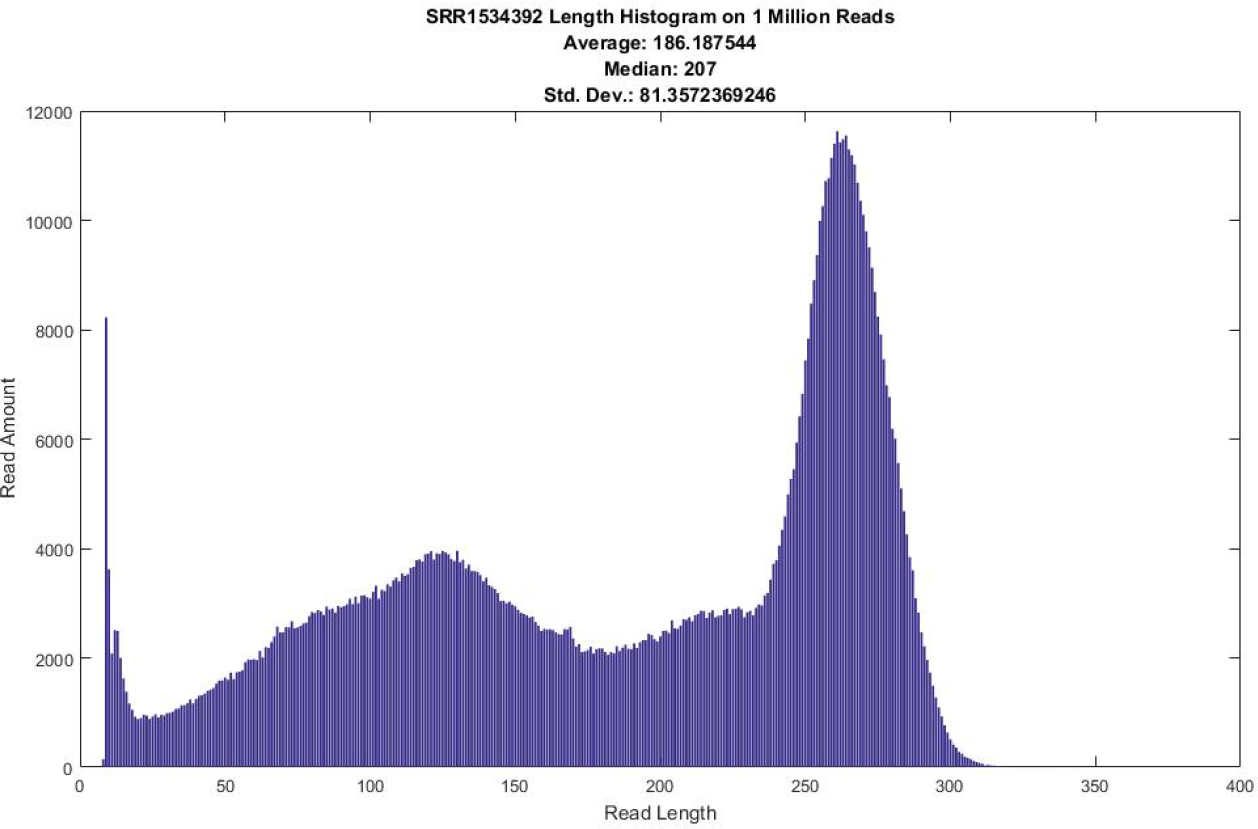
A read length histogram for Ion Torrent reads from data set SRR1534392.

Buckets, each consisting of 500k reads, were created at assorted read lengths from the SRR2842547 and SRR2842546 data, and the uniquely mapped percent was calculated for each mapper with the results visualized in Figure 11. Except for Walt, mapper performance tended to increase with higher read length; however, for BisPin, BWA-Meth, and Tabsat, mapper performance suffered slightly with the longest reads. Bismark had the most dramatic improvement with increasing read length. Both Tabsat and BisPin performed similarily, but BisPin performed the best.

To see if using extra indexes could improve mapper performance for the smallest reads ranging from 9 to 75 base pairs, six indexes of assorted mask lengths with spaced seeds were created and used to align 500k of these reads, but the results were identical to using two indexes. The ineluctable conclusion is that this strategy will have no effect, so improving the indexing strategy was not pursued further.

**Figure 11:**
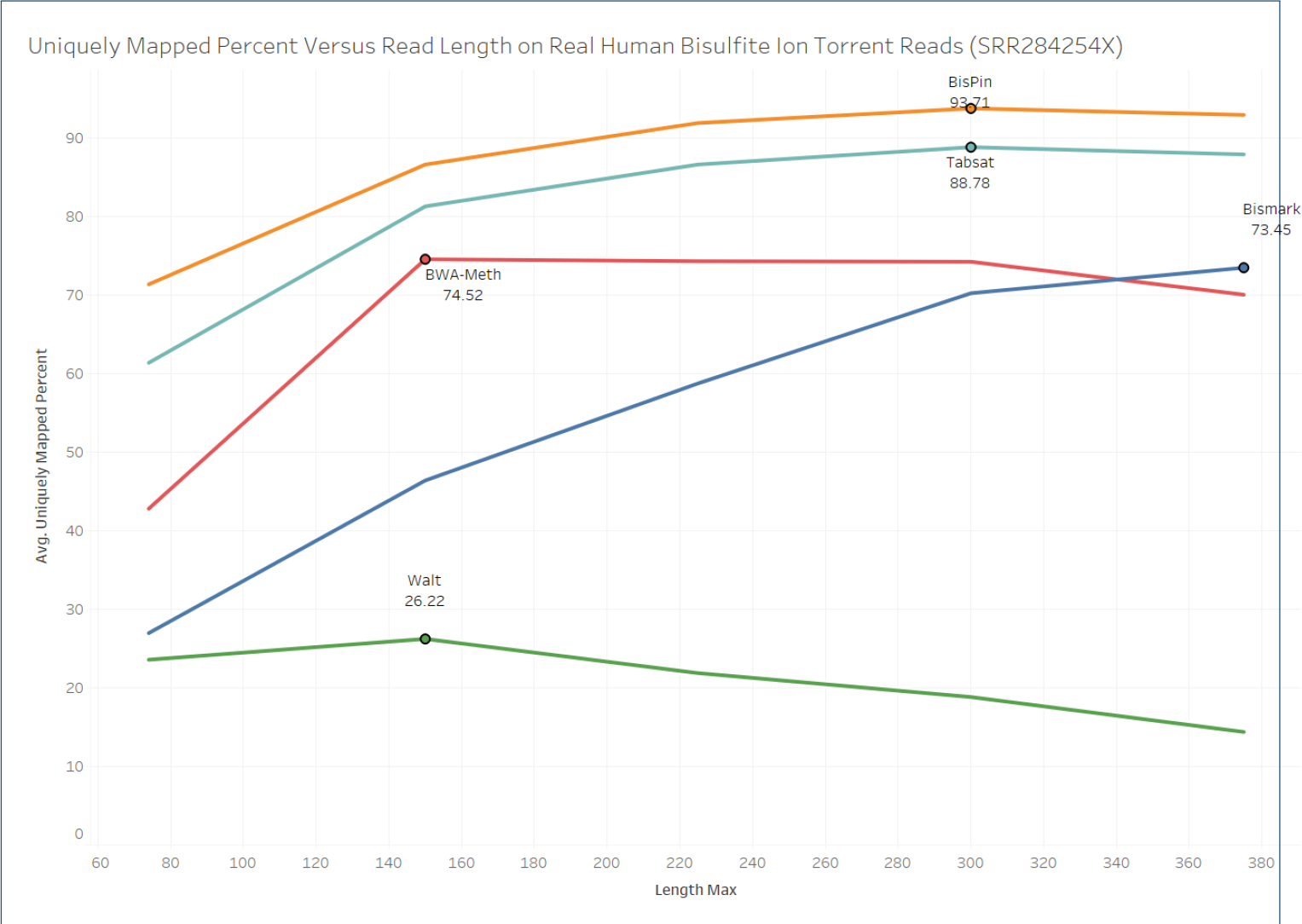
Mapper performance from uniquely mapped percent versus read length on real data. The percent is an average of results from data sets SRR2842547 and SRR2842546.

For the one million bisulfite-treated Ion Torrent mouse SRR1534391 reads, a test of how similar BisPin with BFAST-Gap results were to Tabsat and BWA-Meth was performed by calculating how many reads there were that mapped to the same location on the genome. A read was considered mapped to the same location for two or more read mappers if it was uniquely aligned and if the genome location differed by no more than three bases. The genome location is given in the SAM alignment file in the RNAME and POS fields. A Venn diagram shows the results in Figure 12.

The diagram shows that different read mappers have fairly overlapping results with more than half of the alignments for each read mapper mapping to the same location. This set was 58.758% of BisPin’s uniquely mapped reads, and about one quarter of BisPin’s uniquely mapped reads had a location unique to BisPin. This shows that BisPin’s results are reasonable and similar to other mappers. The set where all read mappers agreed on the location likely has reads correctly mapped. There were 242,833 reads that were uniquely mapped by BFAST-Gap, where other read mappers did not agree on the mapped position. Of these reads, 90% were uniquely mapped to other locations by other read mappers. Because of this and since the locations reported by the read mappers disagreed on approximately 25% — 40% of the reads, this suggests that some kind of ensemble method combining the results of the read mappers could be useful. Perhaps a probabilistic or machine learning model could be employed to combine the results of read mappers.

**Figure 12:**
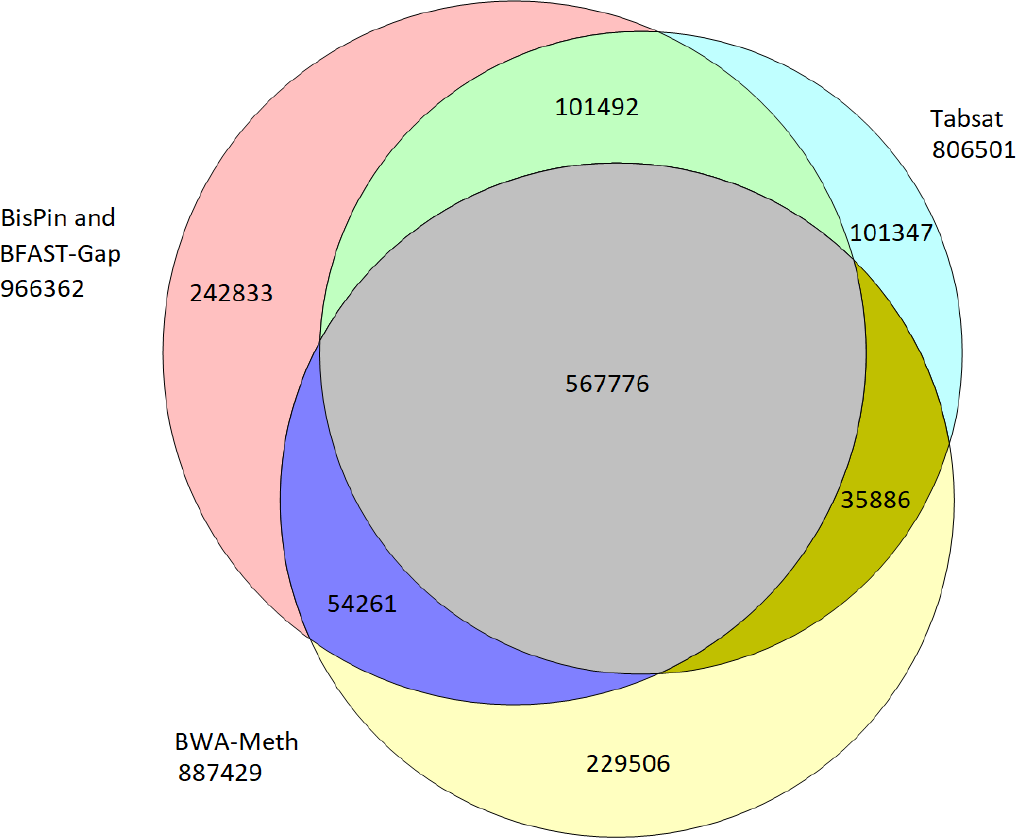
A Venn diagram showing how the uniquely mapped reads from SRR1534391 overlap using mappers BisPin with BFAST-Gap, Tabsat, and BWA-Meth. A read was considered to be in an overlapping set if it was mapped to the same location. The diagram was created with Venn Diagram Plotter available at https://omics.pnl.gov/software/venn-diagram-plotter.

With the tuned logistic gap open function, BisPin with BFAST-Gap was run on one million real and simulated Ion Torrent bisulfite-treated data with no filter. The execution with a logistic gap open penalty and a constant gap extension penalty had slightly more uniquely mapped reads indicating better performance as can be seen in Table 5. Using the logistic function for both gap open and gap extension penalty degraded performance a bit for the simulated data and the SRR1534391 data, but it improved performance by six percent on the SRR2842546 data. Two indexes were used for these executions. This table can be compared directly with Table 4 to compare BisPin using BFAST-Gap with other mappers. BisPin performed the best with this data.

**Table 5:** Logistic gap open and extension penalty function performance on real and simulated data as measured by uniquely mapped percent without a filter. Compare with Table 4.

| Gap Function Type | DWGSIM | SRR1534391 | SRR2842546 |
| --- | --- | --- | --- |
| constant | 96.31859% | 92.25470% | 82.59100% |
| logistic gap open, constant gap extension | 96.46848% | 92.87040% | 83.24190% |
| logistic gap open, logistic gap extension | 95.80231% | 92.4691% | 89.6092% |

Table 6 shows mapper performance on three regular Ion Torrent read data sets. BFAST-Gap did the best with high amounts of uniquely mapped reads on data sets ERR699568 and SRR611141, and it was the second best behind BWA on data SRR2734774. BFAST-Gap has zero reads ambiguously mapped since the homopoly-mer run based gap penalty function tends to make reads uniquely mapped since it uses precise decimal number penalties rather than integer penalties. TMAP did well. Soap2 generally performed the most poorly.

**Table 6:** Regular Ion Torrent mapper performance on three real data sets of one million reads each. The percentage of reads for each category is shown. BFAST-Gap used a tuned logistic function for the gap open and extension penalty functions, and it generally performed well. Soap2 does not distinguish ambiguously mapped reads in its reported statistics.

| Mapper | Unique | Ambiguous | Unmapped or Filtered |
| --- | --- | --- | --- |
| Mouse ERR699568 |  |  |  |
| BFAST-Gap | 91.9665% | 0.0% | 8.0335% |
| TMAP | 86.8102% | 7.95% | 5.2398% |
| Bowtie2 | 64.15% | 33.03% | 2.82% |
| BWA | 86.9847% | 11.1932% | 1.8221% |
| Soap2 | 25.08% | - | 74.92% |
| Human SRR2734774 |  |  |  |
| BFAST-Gap | 88.6083% | 0.0% | 11.3917% |
| TMAP | 88.2288% | 8.8664% | 2.9048% |
| Bowtie2 | 77.55% | 20.21% | 2.23% |
| BWA | 92.4334% | 5.8566% | 1.71% |
| Soap2 | 39.88% | - | 60.12% |
| Human SRR611141 |  |  |  |
| BFAST-Gap | 87.3747% | 0.0% | 12.6253% |
| TMAP | 85.5568% | 9.2704% | 5.1728% |
| Bowtie2 | 75.06% | 21.18% | 3.76% |
| BWA | 86.1396% | 5.9415% | 7.9189% |
| Soap2 | 19.7% | - | 80.3% |

As with bisulfite-treated Ion Torrent reads from Table 4, these mappers are not necessarily tuned for Ion Torrent data, and their performance could be improved by adjusting their parameters. Soap2 can increase the maximum number of mismatches from 5, and can allow a continuous gap sized larger than 0. BWA and Bowtie2 have a minimum score parameter.

### Timing

For alignment time only, on simulated paired-end 250k 100bp reads data, BisPin took 103.5 minutes, Bismark 6.5 minutes, BWA-Meth 3.5 minutes, and Walt 5.5 minutes. BisPin is approximately 20 times slower because BFAST is slow. For one million reads, multiprocessing improved BisPin postprocessing speed, excluding the reference genome load time, by approximately 1.29 times.

On one million reads from the mouse regular Ion Torrent data (SRA#: ERR699568) using 30 threads on a machine with 32 processing cores (E5-2660 @ 2.20GHz with 198 GB of memory), BFAST-Gap took about one hour. Bowtie2 took 5 minutes, and TMAP took 2 minutes. BWA and Soap2 each took approximately half a minute. BFAST-Gap is substantially slower.

## Conclusion

Mapping bisulfite reads is challenging. BisPin employs several strategies to address these difficulties. It uses rescoring to disambiguate some ambiguously mapped reads, and can perform extra processing with additional indexes to attempt to align unmapped reads. It uses multiprocessing and multithreading for improved run-time efficiency, and can align a selected partition of reads from a file for deployment on a compute cluster. BisPin supports a variety of popular read constructions and layouts including the hairpin construction strategy, which is eminently useful for bisulfite-treated reads. BisPin performed well with a variety of real and simulated data compared to other read mappers.

BisPin with BFAST-Gap improves upon read mapping with bisulfite-treated Ion Torrent reads, and BFAST-Gap improves upon read mapping for regular Ion Torrent reads compared to other read mappers including TMAP, which was designed for Ion Torrent reads, in some ways.

For improving run-time, BFAST could be modified to include a highly optimized SIMD implementation of the Smith-Waterman algorithm as in Bowtie2, and it could dispense with intermediate files, which are slow to create. BFAST has parameters that could be adjusted that might improve run-time at the expense of accuracy. An example of this is filtering out reads that match to too many locations.

Perhaps BisPin’s hairpin sequencing approaches could be applied to Oxford Nanopore data because it uses a hairpin connector. Ranking the alignments with a probability measure could be more precise and informative as is done in [42], but it could be less time efficient as computing a probability may require more calculations.

## Author’s contributions

Jacob Porter created BisPin, modified BFAST to produce BFAST-Gap, performed the data analysis, and wrote the paper, and Liqing Zhang consulted on the data analysis and edited the manuscript.

## Acknowledgements

The hairpin data was provided by Hehuang Xie of the Biocomplexity Institute of Virginia Tech. Hong Tran provided the Python software to simulate bisulfite treatment.

## Availability of data and materials

BisPin (Python 2.7) and BFAST-Gap (C++) are available at https://www.github.com/JacobPorter under a GNU General Public License.

